# Impact of population structure in the design of RNA-based diagnostics for antibiotic resistance in *Neisseria gonorrhoeae*

**DOI:** 10.1101/537175

**Authors:** Crista B. Wadsworth, Mohamad R.A. Sater, Roby P. Bhattacharyya, Yonatan H. Grad

**Affiliations:** Department of Immunology and Infectious Diseases, Harvard T.H. Chan School of Public Health, Boston, MA 02115; Infectious Disease and Microbiome Program, Broad Institute of MIT and Harvard, Cambridge, MA 02142; Division of Infectious Diseases and Department of Medicine, Massachusetts General Hospital, Boston, MA 02114; Division of Infectious Diseases, Brigham and Women’s Hospital, Harvard Medical School, Boston, MA 02115

**Author notes:** Corresponding Authors: Crista Wadsworth, Yonatan Grad.

**Keywords:** *Neisseria gonorrhoeae*, gonorrhea, antimicrobial susceptibility testing, AST, macrolide, azithromycin, ciprofloxacin, gene expression, RNA-seq, transcriptome profiling

## Abstract

Quantitative assessment of antibiotic-responsive RNA transcripts holds promise for a rapid point of care (POC) diagnostic tool for antimicrobial susceptibility testing. These assays aim to distinguish susceptible and resistant isolates by transcriptional differences upon drug exposure. However, an often-overlooked dimension of designing these tests is that the genetic diversity within a species may yield differential transcriptional regulation independent of resistance phenotype. Here, we use a phylogenetically diverse panel of *Neisseria gonorrhoeae* and transcriptome profiling coupled with RT-qPCR to test this hypothesis, to identify azithromycin responsive transcripts and evaluate their potential diagnostic value, and to evaluate previously reported diagnostic markers for ciprofloxacin resistance (*porB* and *rpmB*). Transcriptome profiling confirmed evidence of population structure in transcriptional response to azithromycin. Taking this population structure into account, we found azithromycin-responsive transcripts overrepresented in susceptible strains compared to resistant strains, and selected four candidate diagnostic transcripts (*rpsO*, *rplN*, *omp3*, and NGO1079) that were the most significantly differentially regulated between phenotypes across drug exposure. RNA signatures for these markers categorically predicted resistance in 19/20 cases, with the one incorrect categorical assignment for an isolate at the threshold of reduced susceptibility. Finally, we found that *porB* and *rpmB* expression were not diagnostic of ciprofloxacin resistance in a panel of isolates with unbiased phylogenetic sampling. Overall, our results suggest that RNA signatures as a diagnostic tool are promising for future POC diagnostics; however, development and testing should consider representative genetic diversity of the target pathogen.

## INTRODUCTION

The rise of resistance to antimicrobial drugs presents a critical threat to the successful treatment of infectious diseases. Rapid point of care (POC) diagnostics for drug susceptibility could guide therapeutic decisions, thereby improving time to appropriate therapy and limiting the extent of broad empiric antibiotic use. DNA-based susceptibility prediction tests are promising and may have niche applications, but are subject to concerns about the comprehensiveness with which genotypic features accurately predict resistance (i.e., due to complex epistasis or the additive effects of other loci) and are also sensitive to the emergence of novel resistance alleles through mutation or horizontal gene transfer. In contrast, quantitative assessment of antibiotic-responsive RNA transcripts can rapidly differentiate pathogen strains by resistance phenotype independent of resistance mechanism or genetic background. Some of the earliest cellular responses to antibiotic exposure are mediated through transcriptional changes, which has recently been demonstrated in *Escherichia coli*, *Pseudomonas aeruginosa*, *Klebsiella pneumoniae*, *Acinetobacter baumannii*, and *Neisseria gonorrhoeae* (1–4). These RNA-based tests are particularly attractive as POC diagnostics as drug sensitive and resistant bacteria proceed down distinct physiological pathways indicative of cellular stress or normal cell growth in as little as 10 minutes (3) – much faster than the current ‘gold standard’ for resistance phenotyping via minimum inhibitory concentration (MIC) by agar dilution, e-test, or disk diffusion which can take days to months depending on bacterial growth rate.

Consideration of the impact of genetic diversity within the targeted pathogen species will be key to the success of effective RNA-based antimicrobial resistance diagnostics. Gene expression heterogeneity is known to occur between distinct lineages within bacterial species (5–7), and this strain-to-strain variation leaves the open possibility that transcripts may be antibiotic-responsive in some strains but not others of the same phenotype. Thus, to identify antibiotic-responsive transcripts that define the resistance phenotype independent of genetic mechanism of resistance and genomic background, panels of clinical isolates must be surveyed to characterize the full transcriptional diversity of a species. However, while large panels of isolates have been used before to search for diagnostic transcripts (1–4), lack of consideration of phylogenetic structure and specific attention to diverse clade sampling within the assay design panel risks mistakenly attributing distinct RNA signatures to resistance phenotype when they instead derive from population structure (i.e., as a result of genetic drift or divergent selection).

Here, we interrogate the impact of phylogenetic distance on strain variability in gene expression, and search for antibiotic-responsive RNA transcripts diagnostic of antimicrobial resistance in *Neisseria gonorrhoeae,* the Gram-negative pathogen that causes the sexually transmitted disease gonorrhea. Antimicrobial resistance in *N. gonorrhoeae* is one of the most critical contemporary threats to public health with resistance emerging to every class of antimicrobials that have been used to empirically treat gonococcal infection (i.e., penicillins, sulfonamides, tetracyclines, fluoroquinolones, macrolides, and cephalosporins) (8, 9). With only a few novel antimicrobials and combination therapies currently in development (10–12) and rapidly rising incidence of infection (13), we face an imminent threat of untreatable gonorrhea (9, 14). POC tests that move healthcare providers away from empiric treatment to data-driven prescription are urgently needed to support good antimicrobial stewardship by rapidly dispensing effective therapeutics to individuals and their sexual contacts and minimizing the selective pressures on bacterial bystanders from inappropriate treatment (15).

We focused on RNA-based diagnostic assay development for two clinically relevant drugs: azithromycin and ciprofloxacin. Azithromycin is a macrolide antibiotic that inhibits protein synthesis by binding to the 23S rRNA component of the 50S ribosome, and is one of the two first-line drugs recommended as dual-antimicrobial therapy for uncomplicated cases of gonococcal infection by the U.S. Centers for Disease Control and Prevention (CDC) (16). Azithromycin resistance is especially attractive for phenotypic diagnostic development, as there are multiple resistance mechanisms, including as yet unexplained pathways (17, 18). Resistance mechanisms that have been associated with or experimentally confirmed to be involved in reduced susceptibility to azithromycin include: mutations in the 23S rRNA azithromycin binding sites (C2611T and A2059G) (17, 19, 20), mosaic multiple transferable resistance (*mtr*) efflux pump alleles acquired from commensal *Neisseria* species (17, 18, 21–23), mutations that enhance the expression of Mtr (24–26), mutations in *rplD* (17), *rplV* tandem duplications (17), and variants of the rRNA methylase genes *ermC* and *ermB* (27). Ciprofloxacin is a quinolone antibiotic that targets DNA gyrase (encoded by *gyrA*) and topoisomerase IV (encoded by *parC*), and was recommended as a first-line therapy for gonorrhea until 2007, at which point the prevalence of quinolone resistance overtook the 5% accepted threshold for suggested use (28). However, the development of a DNA-based diagnostic for quinolone susceptibility has led to the introduction of targeted quinolone use (29), and as such makes an attractive target for rapid POC diagnostics.

Here, we used RNA-seq to profile the transcriptomes of a panel of *N. gonorrhoeae* isolates that are representative of the known diversity within the gonococcal population (17), in response to exposure to azithromycin. Using the transcriptomes from this panel, we ask: is there a differential transcriptome-wide response to azithromycin between resistant and susceptible isolates, what is the impact of evolutionary distance on transcriptome regulation, and are RNA signatures of particular transcripts ubiquitously diagnostic of resistance phenotype across the species? After defining candidate diagnostic markers for azithromycin resistance, we then verify them via RT-qPCR in an expanded panel of isolates. We also assess the previously described diagnostic markers for ciprofloxacin resistance in *N. gonorrhoeae*, *porB* and *rpmB* (3), in a phylogenetically representative panel of isolates.

## RESULTS

### Transcription profiling

Whole transcriptome profiling was used to identify differentially expressed azithromycin-responsive RNA transcripts between resistant and susceptible isolates of *N. gonorrhoeae* after 60 and 180 minutes of below breakpoint azithromycin exposure (0.125 μg/ml; Figure 1). Clinical isolates were sampled from across the known phylogenetic diversity of the species, based on the collection reported in Grad et al. (17). This included paired resistant (MIC ≥ 1 μg/ml) and susceptible (MIC < 1 μg/ml) strains from four of the twelve sub-structured clades, termed Bayesian analysis of population structure (BAPS) groups (17) (Table 1; Figure 2A). In total ten isolates were selected: four susceptible isolates, two resistant isolates with mosaic *mtr* efflux pump alleles acquired from commensal *Neisseria*, three resistant isolates with the C2611T 23s rRNA mutation, and one resistant isolate with the A2059G 23s rRNA mutation (17, 18). A total of ~258 million 50 bp paired-end reads were generated across 90 libraries. Each library had on average 3.17 ± 0.16 million reads, and an average of 1.82 ± 0.11 million reads per library mapped to the FA1090 (AE004969.1) reference genome.

**Table 1.** Properties of the clinical isolates used to develop diagnostic markers for azithromycin resistance (n=20)

| Isolate | Location | Collection Year | BAPS | AZI MIC (ug/ml) | Resistance Category | Resistance Mechanism |
| --- | --- | --- | --- | --- | --- | --- |
| **Whole Transcriptome** |  |  |  |  |  |  |
| GCGS0524 | Albuquerque, NM | 2000 | 4 | 0.25 | Susceptible | - |
| GCGS0276 | Kansas City, MO | 2000 | 4 | 1 | Resistant | *mtr* mosaic |
| GCGS0745 | Honolulu, HI | 2012 | 4 | 256 | Resistant | 23S A2059G |
| GCGS0996 | Seattle, WA | 2012 | 6 | 0.5 | Susceptible | - |
| GCGS0834 | Los Angeles, CA | 2012 | 6 | 2 | Resistant | *mtr* mosaic |
| GCGS0641 | Orange County, CA | 2011 | 7 | 0.25 | Susceptible | - |
| GCGS0354 | Dallas, TX | 2012 | 7 | 8 | Resistant | 23S C2599T |
| GCGS0550 | Birmingham, AL | 2012 | 7 | 8 | Resistant | 23S C2599T |
| GCGS0353 | Dallas, TX | 2011 | 8 | 0.03 | Susceptible | - |
| GCGS0363 | Detroit, MI | 2011 | 8 | 4 | Resistant | 23S C2599T |
| **RT-qPCR** |  |  |  |  |  |  |
| GCGS1035 | Detroit, MI | 2007 | 1 | 0.5 | Susceptible | - |
| GCGS0481 | Portland, OR | 2006 | 1 | 2 | Resistant | *mtr* mosaic |
| GCGS0926 | Minneapolis, MA | 2012 | 2 | 0.125 | Susceptible | - |
| GCGS0298 | San Diego, CA | 2005 | 2 | 2 | Resistant | *mtr* mosaic |
| GCGS0605 | Miami, FL | 2002 | 3 | 0.25 | Susceptible | - |
| GCGS1026 | Chicago, IL | 2008 | 5 | 16 | Resistant | *rpIV* tandem duplication |
| GCGS0436 | Orange County, CA | 2010 | 7 | 2 | Resistant | 23S C2599T |
| GCGS0465 | Pheonix, AZ | 2011 | 10 | 0.06 | Susceptible | - |
| GCGS0525 | Atlanta, GA | 2000 | 11 | 2 | Resistant | *mtr* mosaic |
| GCGS0313 | Balitimore, PA | 2012 | 12 | 0.06 | Susceptible | - |

**Figure 1.**
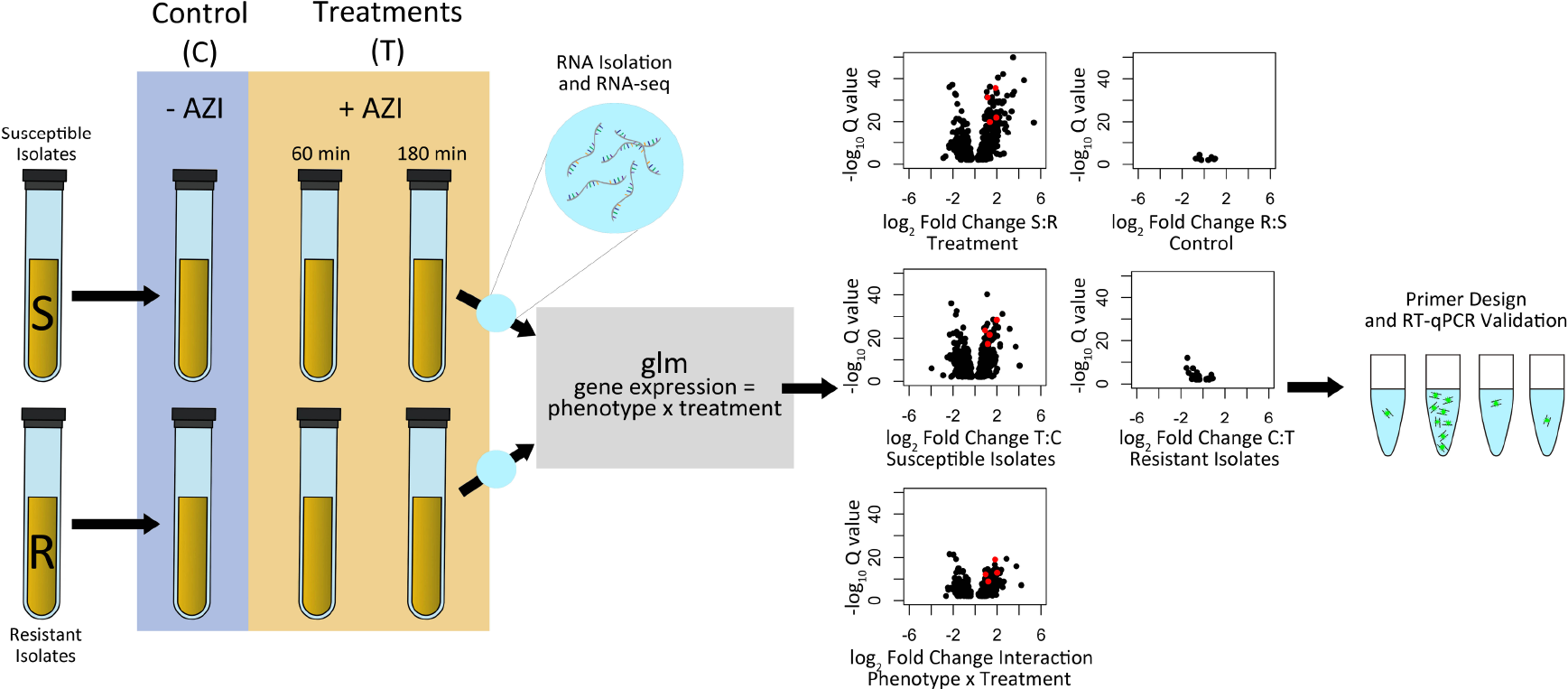
Workflow for generating candidate diagnostic markers for azithromycin resistance. We sampled cultures of susceptible (MIC < 1 μg/ml) and resistant (MIC ≥1 μg/ml) isolates prior to the addition of azithromycin, after 60 minutes of exposure to 0.125 μg/ml of azithromycin, and after 180 minutes of exposure to 0.125 μg/ml of azithromycin. RNA-seq libraries were then constructed and sequenced; and a generalized linear model was fit to the resultant data with phenotype (resistant *vs.* susceptible), treatment (control *vs.* later time points), and the interaction between phenotype and treatment used as model terms. Volcano plots are shown for the control to 60-minute exposure comparisons with a FDR cutoff of ≤ 0.01. Black circles indicate all significant transcripts for each model contrast, and red dots indicate the four candidate diagnostic markers ultimately chosen for RT-qPCR validation. Most of the observed response occurred (1) in susceptible strains between the control and the drug exposed conditions, (2) between resistant and susceptible isolates in the antibiotic exposure condition, and (3) having a significant interaction term indicating differential response between susceptible and resistance isolates across treatments.

**Figure 2.**
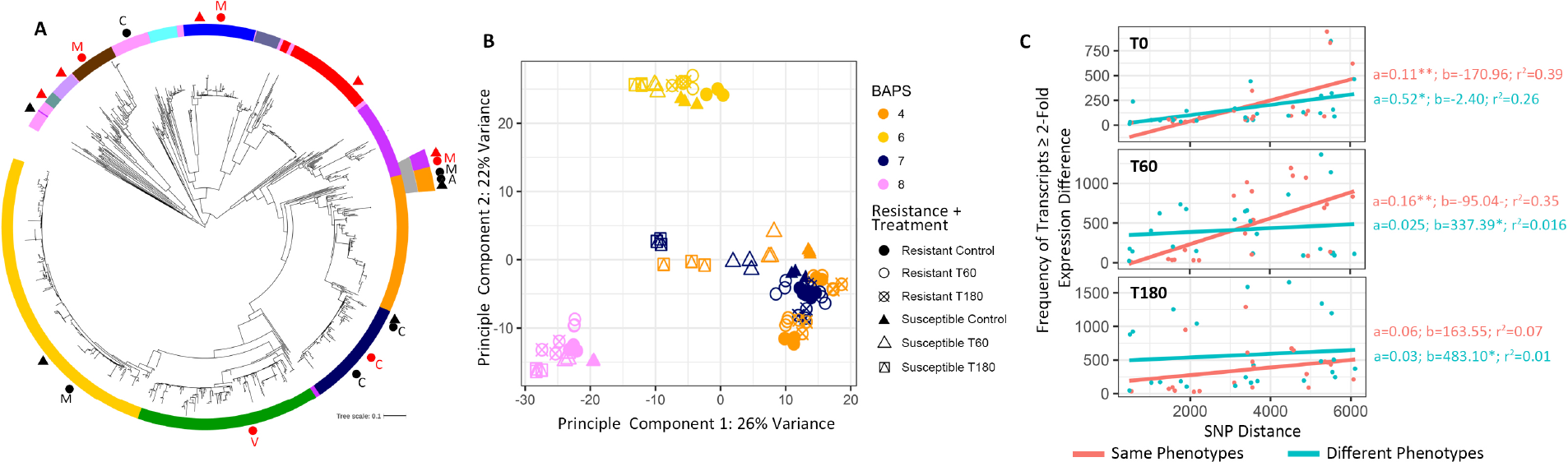
Genetic distance impacts transcriptional regulation in ten phylogenetically diverse isolates. (A) A maximum likelihood whole-genome-sequence phylogeny of 1102 *Neisseria gonorrhoeae* isolates from GISP, based on single-nucleotide polymorphisms generated from mapping to the FA1090 reference genome (Grad et al. 2016). The annotation ring displays the Bayesian analysis of population structure (BAPS) groups. All annotated shapes represent the panel of isolates selected for identifying azithromycin responsive RNA transcripts and evaluating their potential diagnostic value. Black circles represent resistant isolates selected for RNA-seq and RT-qPCR validation, black triangles represent susceptible isolates selected for RNA-seq and RT-qPCR validation, red circles show resistant isolates selected for RT-qPCR validation, and red triangles show susceptible isolates selected for RT-qPCR validation. Resistance mechanisms are indicated by A for the A2059G 23s rRNA mutation, C for the C2611T 23s rRNA mutation, M for the mosaic *mtr* mutation, and V for the *rplV* tandem duplication. (B) For all 90 RNA-seq libraries generated within this study, principal component analysis revealed that the largest variance in gene expression can be attributed to genetic distance between isolates rather than drug exposure, resistance phenotype, or resistance mechanism. (C) We further tested the impact of evolutionary distance on transcriptional regulation, by comparing the number of pairwise SNP differences with the frequency of transcripts with a ≥2 fold expression difference across the ten isolates for which we had collected RNA-seq data at baseline (T0, no drug exposure), after 60 minutes of drug exposure (T60), and after 180 minutes of drug exposure (T180). Linear models were fit to either paired isolate comparisons that belonged to the same (resistant vs. resistant or susceptible vs. susceptible) or different (resistant vs. susceptible) phenotypic classes. In all cases, we observed positive slope values.

### Genetic distance and population structure impact transcriptome regulation

First, we investigated the relationship between phylogenetic structure and transcriptome regulation. Principal component analysis (PCA) revealed three distinct transcriptional clusters along the first two dimensions (PC1 and PC2), representing the greatest variability within the gene expression dataset, which were primarily structured by genetic distance rather than resistance phenotype, resistance mechanism, or drug exposure condition (Figure 2A; Figure 2B). Isolates that belonged to BAPS groups 4 and 7, which were the most genetically similar as indicated by their grouping in adjacent clades, were also the most transcriptionally similar and clustered on the PCA plot (Figure 2A; Figure 2B). These two clades, in addition to containing susceptible isolates, harbored multiple mechanisms of resistance among resistant isolates including: the 23s rRNA C2611T, 23s rRNA A2059G, and mosaic *mtr* efflux pump mutations. Isolates from BAPS groups 6 and 8 clustered independently from each other as well as the other clades sampled, and were also the most evolutionarily distant from other clades on the gonococcal tree (Figure 2A; Figure 2B).

The relationship between genetic distance and transcriptional regulation was also supported by a pairwise comparison of whole-genome SNP distance versus pairwise fold change differences in gene expression across all ten isolates transcriptionally profiled in multiple treatment conditions. SNP distances for all isolate pairs ranged from 488 to 6,106 (mean=3,410). The type of each pairwise comparison was defined as being of either the same (resistant versus resistant or susceptible versus susceptible) or different (resistant versus susceptible) phenotypic classes. For both categories, paired isolates of increasing core genome SNP distances had an increasing number of differentially regulated transcripts (≥ 2-fold expression difference) between them at baseline (T0) and in treatment conditions (T60 and T180), as indicated by each linear model’s positive slope values (Figure 2C). This pattern also held true for all transcripts regardless of their fold change in abundance between isolates (Supplementary Figure 1). Interestingly, at baseline (T0), the two markers previously reported as being diagnostic of ciprofloxacin resistance (3), *porB* and *rpmB*, diverged transcriptionally with increasing evolutionary distance between isolates as well (Supplementary Figure 1).

### Transcriptional response to azithromycin and candidate marker selection

We next investigated the transcriptome-wide response to azithromycin across 60 and 180-minute drug exposure treatments in our phylogenetically diverse isolate panel. To define the differential response patterns of resistant and susceptible isolates across treatment conditions, we fit a generalized linear model to the count data generated from all 90 RNA-seq libraries with phenotype (resistant versus susceptible), treatment (control versus later time points), and the interaction between phenotype and treatment used as model terms. Overall, 978 genes were significantly differentially expressed in at least one comparison involving resistance phenotype or condition from 0 to 60 minutes, and 1,177 genes were significantly differentially expressed from 0 to 180 minutes (Figure 1; Supplementary Figure 2). For both drug exposure time points, the majority of response to condition was initiated only in susceptible isolates (Figure 1; Supplementary Figure 2).

For susceptible isolates, we observed large-scale changes in gene expression in response to azithromycin in both treatments, thus the 60-minute time point was selected for designing candidate diagnostic markers. Candidate markers were required to be significantly differentially expressed between the control and drug exposure in the susceptible strains and between resistant and susceptible isolates under drug exposure and also to have a significant interaction term. From the 413 transcripts that fit these conditions, we selected those that were within the lists of the top 10% of most significant FDR-corrected *P*-values for each comparison to assess as markers via RT-qPCR (Supplementary Figure 2; Supplementary Figure 3).

## RT-qPCR Validation

### Azithromycin

We assessed the diagnostic potential of RNA-seq nominated markers in a large panel of genetically diverse isolates with non-biased phylogenetic sampling using RT-qPCR. In addition to the panel of isolates used for RNA-seq, ten more isolates were chosen from eight of the twelve BAPS groups for validation (Table 1; Figure 2A). Of these, five were susceptible, three were resistant and had mosaic *mtr* alleles, one was resistant and had the C2611T 23s rRNA mutation, and one was resistant and had a tandem duplication in *rplV* that has previously been associated with resistance (17).

Fold-change across control and 60-minute azithromycin exposure treatments was evaluated for four markers (*rpsO*, NGO1079*, omp3,* and *rplN)* using the 2^−ΔΔCT^ method (30, 31), normalizing to an empirically defined control gene (NGO1935) that did not significantly change in expression between any of our RNA-seq *glm* model contrasts (Supplementary Figure 3). Expression change was then compared between resistant and susceptible isolates. For both *rpsO* and NGO1079, change in RNA abundance was significantly different between resistant and susceptible isolates across treatments and there was no overlap in 2^−ΔΔCT^ values between groups (Figure 3A and 3B). While 2^−ΔΔCT^ values for *omp3* and *rplN* were also significantly different between resistant and susceptible isolates, isolates with intermediate MIC values bounding the reduced susceptibly cutoff of 1 μg/ml displayed some overlap between phenotypes (Figure 3C and 3D).

**Figure 3.**
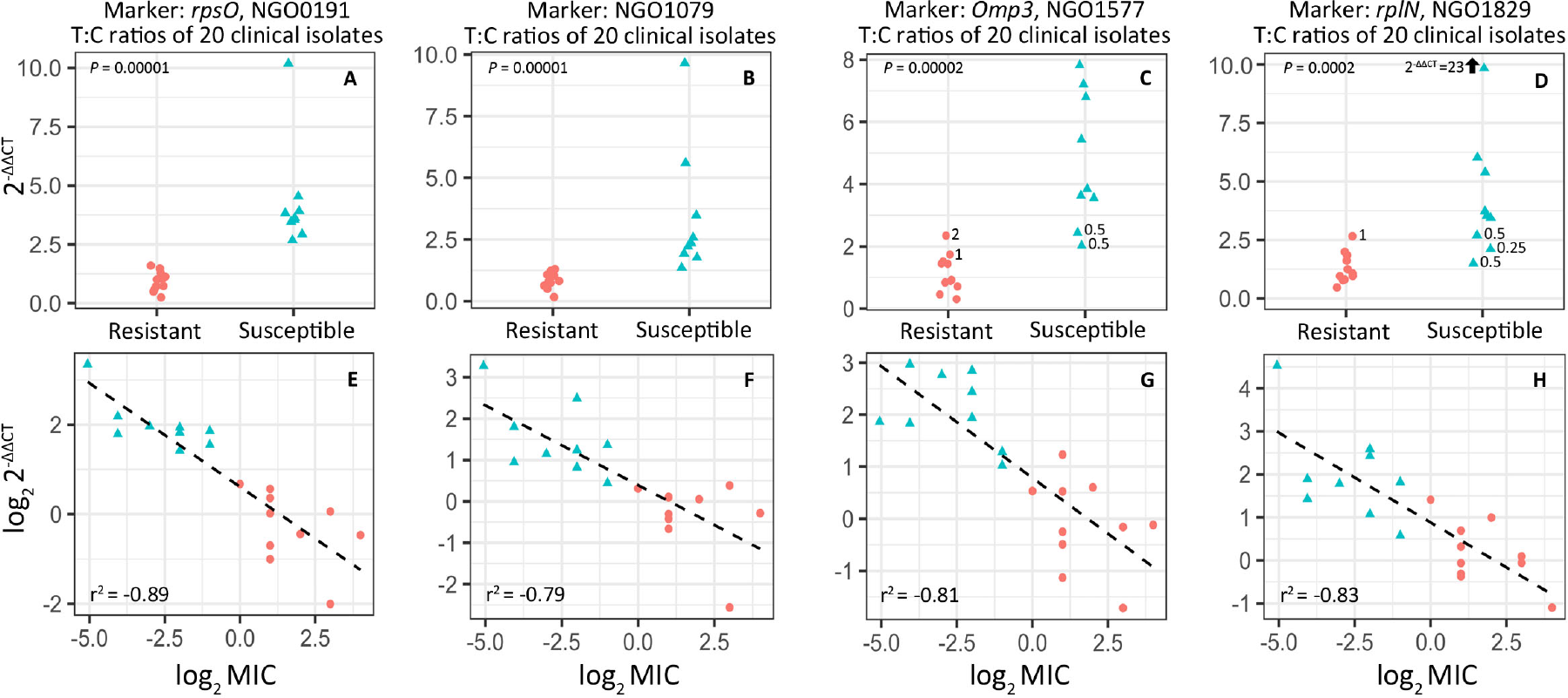
Candidate diagnostic marker validation via RT-qPCR in a panel of 20 clinical isolates (9 susceptible and 11 resistant). (A) NGO0191 or *rpsO*, (B) NGO1079, (C) NGO1577 or *omp3*, and (D) NGO1829 or *rplN* fold change from prior to the addition of azithromycin and 60 minutes after exposure (n=3-4 biological replicates) was calculated using the 2^−ΔΔCT^ method, normalizing to an empirically defined control transcript (NGO1935, *etfB*). All four markers showed significant differences in expression across conditions between resistant and susceptible isolate phenotypic classes, however fold change for *omp3* and *rplN* did overlap between phenotypes for isolates with azithromycin MIC values close to the reduced susceptibility threshold (i.e., 0.25, 0.5, 1, and 2 μg/ml). (E-H) Linear models were fit to log transformed 2^−ΔΔCT^ and MIC values for each marker for each isolate, omitting the GCGS0745 outlier (MIC > 256 μg/ml), and were used to assess the accuracy of 2^−ΔΔCT^ values as a predictor of MIC (Table 2).

We investigated the relationship between the transcriptional response of these markers to azithromycin exposure and MIC by fitting a linear model to the log_2_-transformed 2^−ΔΔCT^ and MIC values, omitting the GCGS0745 outlier isolate (MIC > 256 μg/ml). For all markers there was a strong negative relationship between these two variables, with r^2^ values ranging between −0.79 and −0.89 (Figure 3E-H). Fitted models were subsequently used to estimate the MICs of each isolate given 2^−ΔΔCT^ for each marker, which were then averaged across markers, and rounded to the nearest MIC on the discreet scale used by the Center for Disease Control and Prevention’s Gonococcal Isolate Surveillance Project (GISP) clinical microbiology laboratories (32) (Table 2). Using this method, RNA signatures for these markers were able to categorically predict resistance in 19/20 cases, with the one susceptible isolate called as resistant predicted to be within 1 dilution of the actual MIC (predicted at 1 μg/ml, phenotyped at 0.5 μg/ml), and on the cusp of the reduced susceptibility phenotypic designation. Using this method, we could also predict the MIC of each isolate within + or − 2 dilutions of the tested MIC in 18/20 of cases. For the two isolates for which MICs were incorrectly assigned, the resistance category was called correctly.

**Table 2.**
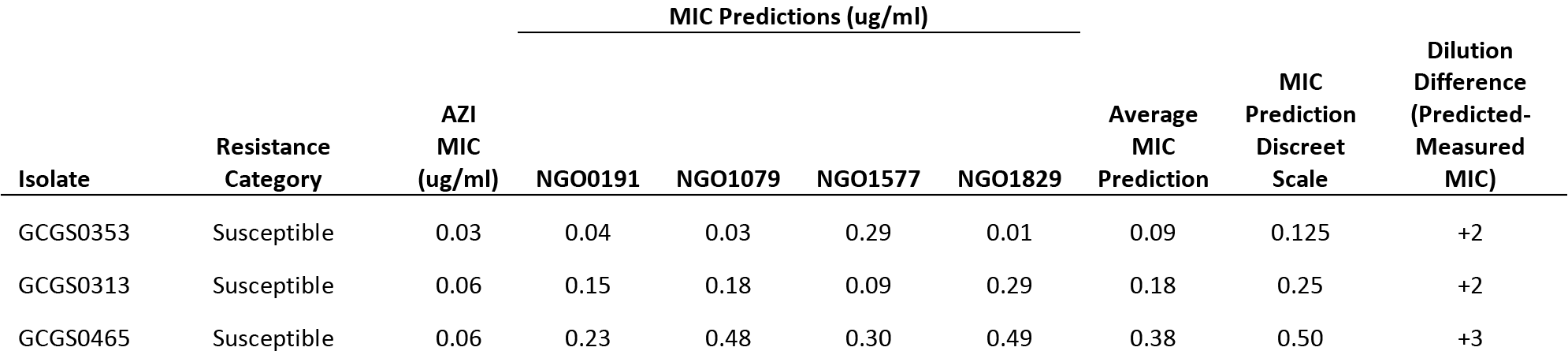
RNA response to 60 minutes of azithromycin exposure (∆∆CT) as a predictor of minimum inhibitory concentration.

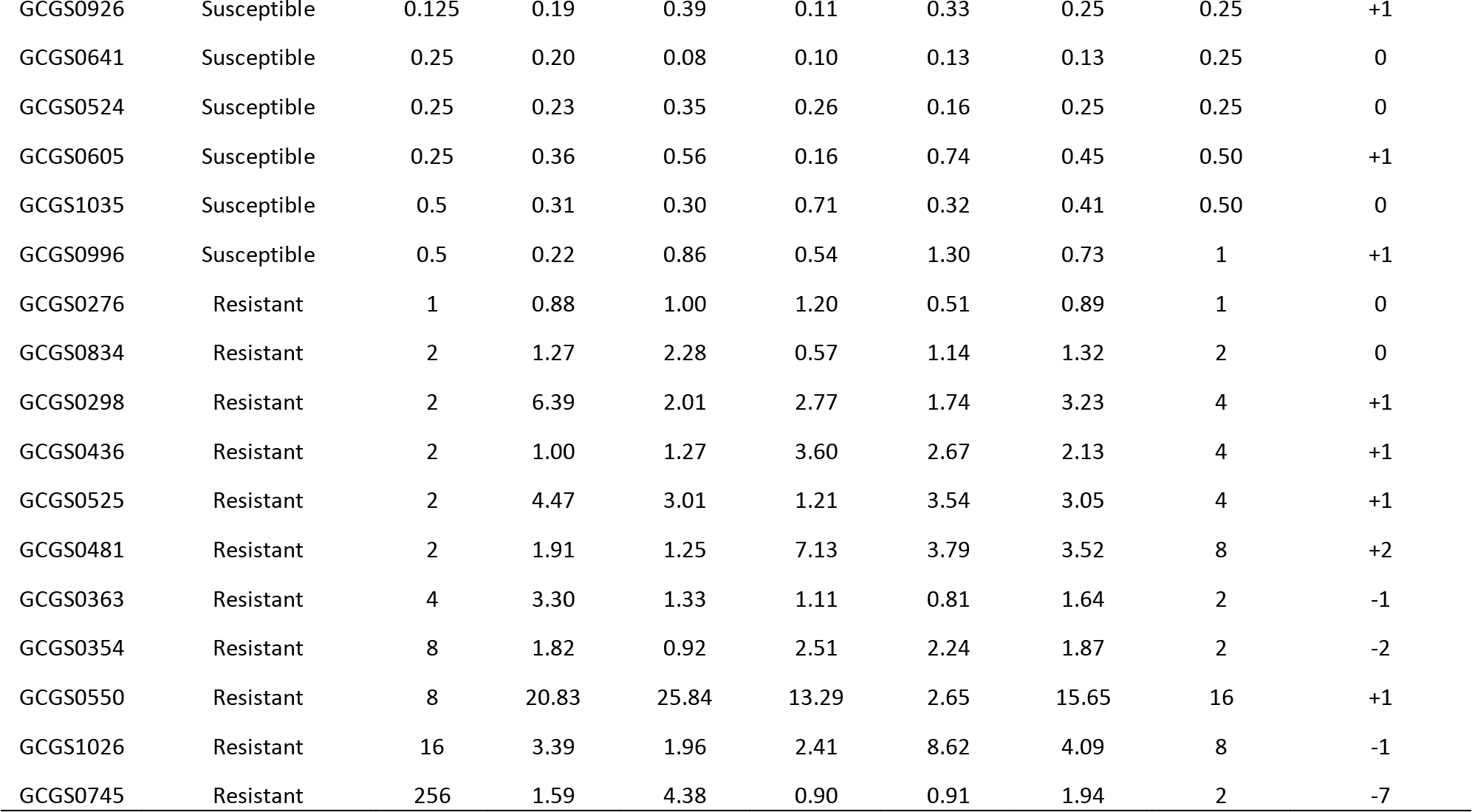

### Ciprofloxacin

For our ciprofloxacin test panel, we selected a set of isolates from the panel that had previously been used to develop diagnostic markers for ciprofloxacin resistance (3) and also added isolates that represented a more phylogenetically comprehensive sampling. The set of isolates used for diagnostic marker development included six resistant isolates (MIC ≥ 1 μg/ml) with *parC* (S87R) and *gyrA* (S91F, D95G) mutations from a single BAPS clade, and five susceptible isolates (MIC < 1 μg/ml) from BAPS clades located elsewhere on the tree (Table 3; Figure 4A). The additional isolates tested here included two resistant isolates with unknown mechanisms of resistance; two resistant isolates with the ParC S87R, GyrA S91F/D95G haplotype; one resistant isolate with the ParC S87R, GyrA S91F/D95A haplotype; and seven susceptible isolates (Table 3; Figure 4A).

**Table 3.**
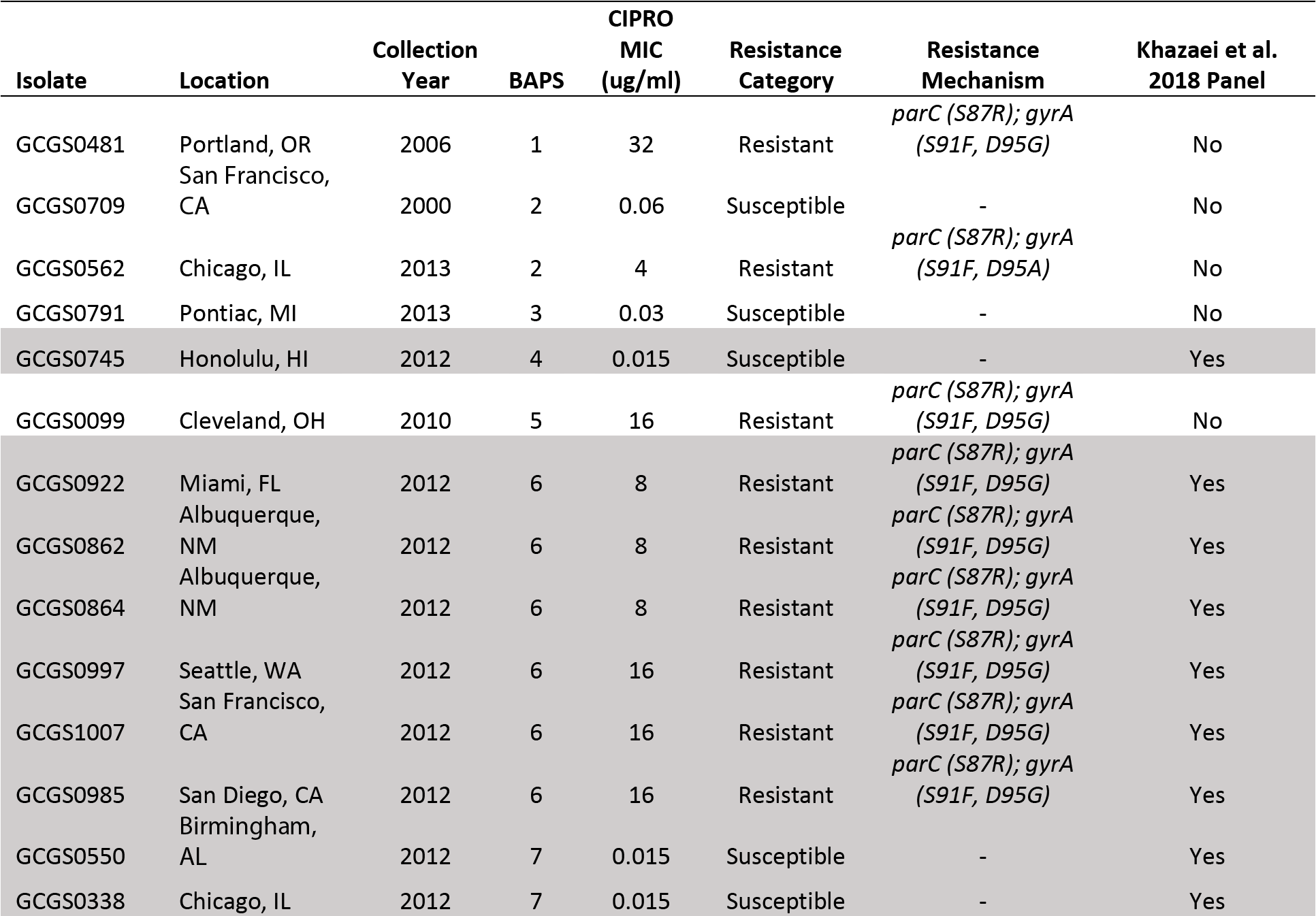
Properties of the clinical isolates used to validate diagnostic markers for ciprofloxacin resistance (n=23)

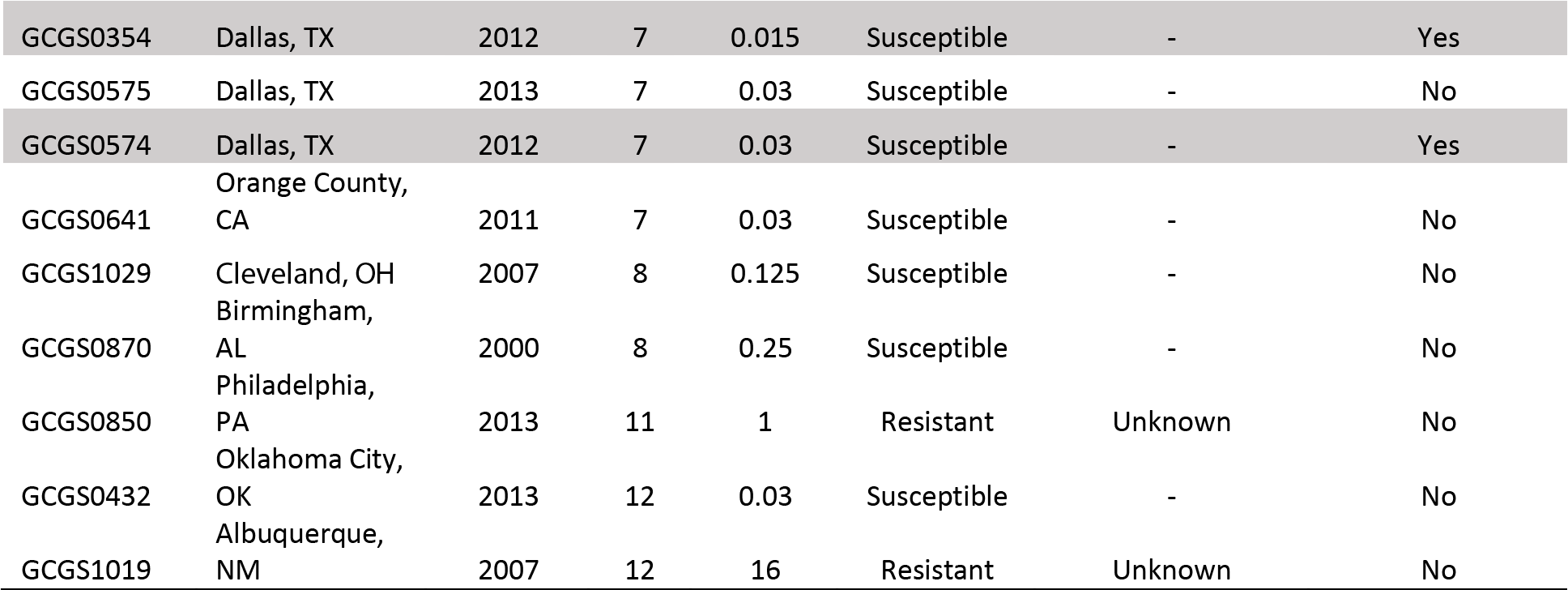

**Figure 4.**
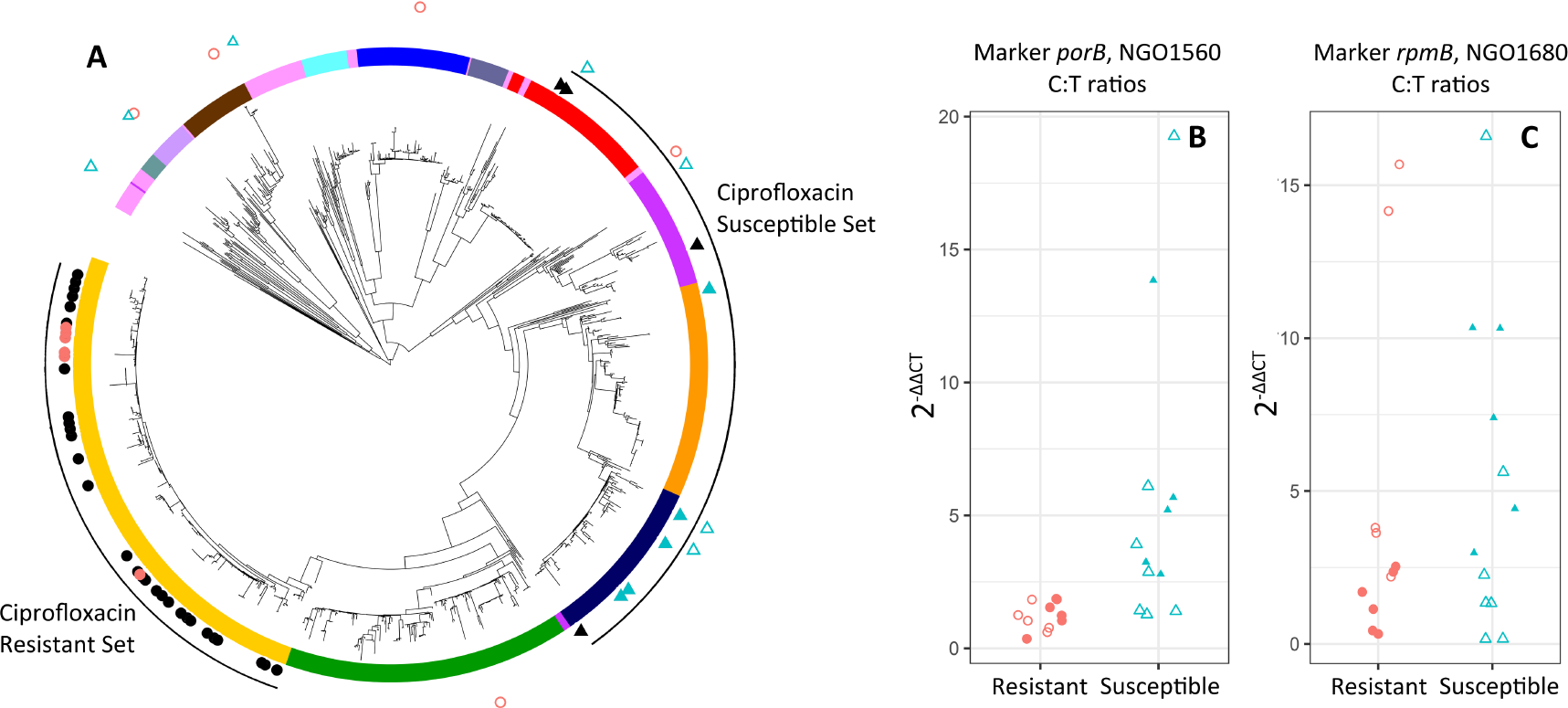
Phylogenetic structure obscures diagnostic signatures of ciprofloxacin resistance. (A) The panel of isolates used to nominate *porB* and *rpmB* as diagnostic phenotypic markers for ciprofloxacin resistance had a highly structured distribution within the gonococcal population, with resistant isolates all sampled from a single clade (closed circles) and susceptible isolates sampled from distantly related clades (closed triangles). In this study, we sample a subset of the panel previously evaluated (3) (closed red circles and blue triangles), in addition to a panel of diverse resistant and susceptible isolates with a less biased sampling pattern (open red circles and blue triangles). RT-qPCR validation revealed the nominated markers *porB* (B) and *rpmB* (C) were diagnostic of resistance in the previously evaluated subset (closed red circles and blue triangles do not overlap); however, in the expanded panel the markers were not diagnostic (open red circles and blue triangles overlap).

Evaluation of the full panel of isolates used by Khazaei et al. (3) within the known genetic diversity of the species revealed a clade-biased ciprofloxacin susceptibility sampling pattern, with resistant isolates all belonging to a single clade and susceptible isolates sampled from elsewhere on the phylogeny (Figure 4A). Both *porB* and *rpmB* were diagnostic of reduced susceptibility to ciprofloxacin in the sample of isolates used by Khazaei et al. (3) that we tested, with no overlap in 2^−ΔΔCT^ values (Figure 4B,C). However, these markers displayed divergent transcription as a function of increasing genetic distance (Supplementary Figure 1), and in the expanded panel of isolates, including a more representative sampling of the known gonococcal genetic diversity, 2^−ΔΔCT^ values of *porB* and *rpmB* were not diagnostic of ciprofloxacin resistance phenotype (Figure 4B,C)

## DISCUSSION

RNA-based resistance diagnostics have the potential to revolutionize antimicrobial selection at the point of care. However, the design of diagnostic tests that rely on changes in RNA-abundance after drug exposure can be vulnerable to erroneous candidate marker designation due to the potential for gene regulation divergence to reflect genetic distance and population structure rather than resistance phenotype. This danger can be amplified if the sampling of isolates for marker design is highly phylogenetically structured by resistance phenotype. Here, we used a combination of whole transcriptome profiling and RT-qPCR to test the importance of controlling for genetic diversity in the design of RNA-based tests, to identify azithromycin responsive transcripts and evaluate their potential diagnostic value, and to evaluate previously reported diagnostic markers for ciprofloxacin resistance (*porB* and *rpmB*).

We expected that if genetic distance was the main contributor to transcriptional variation between isolates, increasing genetic distance would also yield increasing differences in transcriptome regulation, regardless of the resistance phenotypes of the isolates being compared (see models in Supplementary Figure 5). Results from our principal component analysis demonstrated that for our panel of diverse gonococcal isolates, divergence in transcriptional profiles primarily reflected population structure, as indicated by the clustering of isolates along PC1 and PC2 by phylogenetic proximity (i.e., genetic relatedness) as opposed to resistance phenotype, resistance mutations present (mosaic *mtr* or 23s rRNA), or drug exposure condition (Figure 2B).

To further investigate the importance of controlling for phenotype-specific phylogenetic structure in assay design panels, we quantified the pairwise inter-strain whole-genome SNP distances, as a metric of evolutionary distance, and pairwise inter-strain frequency of 2-fold or greater expression differences by transcript across drug exposure conditions, as a metric of transcriptional regulatory evolution (Figure 2C). Linear models fit to either same- or cross-phenotype comparisons indicated several conclusions. First, at baseline for both comparison types, regressions had significant and positive slopes, and y-intercepts that were not significantly different than zero (Figure 2C T0), supporting genetic distance as the main contributor to gene expression variance between isolates in this condition. Second, after 60 minutes of azithromycin exposure, though genetic distance remained the main contributor to gene expression variance in the same phenotypic class comparison, for isolates of different phenotypic classes the slope was no longer significantly different than zero and the y-intercept of 337.39 was significantly greater than zero. This suggests that differences in resistance phenotype had a greater impact on gene expression divergence in this condition than genetic distance (Figure 2C T60). Third, after 180 minutes of azithromycin exposure for isolates of the different phenotypic class comparison the slope was again not significantly different than zero and the y-intercept of 483.10 was significantly greater than zero (Figure 2C T180). In contrast to our principal component analysis supporting the importance of controlling for phenotype-specific clade sampling bias in the assay design panel, these results suggest that gene expression variance driven by differences in phenotype will overwhelm any genetic distance effects, as isolates of different resistance phenotypes are diverted along alternative physiological trajectories in response to drug exposure. However, we suggest a measure of caution in this interpretation, as positive slope values in every treatment condition for cross-phenotype comparisons indicate that some variance in gene regulation is still likely driven by genetic distance effects rather than resistance phenotype alone.

To identify candidate diagnostic markers for azithromycin resistance, we broadly sampled isolates from the *N. gonorrhoeae* phylogenetic tree and selected four candidate diagnostic markers (*rpsO*, *rplN*, *omp3*, and NGO1079). For all candidate transcripts, RNA abundance increased after 60 minutes of azithromycin exposure in susceptible but not resistant isolates (Fig 3), and upregulation was consistent with gene expression results from previous drug challenge studies. Macrolides induce translational stalling by binding the ribosome peptide exit tunnel and inhibiting the exit of nascent proteins, which has been shown to induce the upregulation of ribosomal protein-encoding transcripts (33–35). Upregulation of the 30S and 50S ribosomal protein-encoding transcripts *rpsO* and *rplN* in susceptible isolates of *N. gonorrhoeae* therefore likely serves as a good phenotypic signature of susceptibility. The *omp3* gene encodes the major gonococcal outer-membrane protein Rmp, which (1) has been implicated in the formation, stabilization, and operation of the trimeric porin structure of PorB; and (2) contains a peptidoglycan-binding motif, suggesting a role in crosslinking the outer-membrane proteins to the peptidoglycan and the maintenance of cell membrane integrity (36, 37). Interestingly, the Rmp protein is also upregulated after spectinomycin exposure in susceptible *N. gonorrhoeae* isolates (38). Finally, the putative oxidoreductase NGO1079 was upregulated upon drug exposure in our study. NGO1079 has previously been shown to be upregulated by the AraC-like regulator MpeR, and the expression of MpeR also indirectly modulates resistance to macrolides through transcriptional suppression of the repressor of the *mtrCDE* efflux pump (39). Though all transcripts were significantly upregulated in susceptible isolates compared to resistant isolates across treatment conditions, isolates with intermediate MIC values bounding the reduced susceptibly threshold had some overlap in 2^−ΔΔCT^ values between phenotypic groups for some markers (Figure 3C and 3D). For all tested markers we uncovered a strong negative relationship between the log_2_-transformed 2^−ΔΔCT^ and MIC values (Figure 3E-H). Using regression of these two variables across all tested markers, we accurately predicted resistance category in 19/20 cases (Table 2), with the one susceptible isolate called as resistant on the boarder of the reduced susceptibility phenotypic designation (predicted at 1 μg/ml, phenotyped at 0.5 μg/ml). Though 2^−ΔΔCT^ values could not perfectly predict resistance for isolates with intermediate MICs, results are promising and continued optimization of the assay conditions (markers, drug concentration, exposure time, etc.) may increase diagnostic accuracy.

The sampling scheme for the assay design panel isolates used to identify the transcripts of *porB* and *rpmB* as diagnostic for ciprofloxacin resistance (3) had a clade bias, with all resistant isolates from a single clade and all susceptible isolates from outside this clade (Fig 4A). Additionally, transcription of *porB* and *rpmB* was clearly impacted by evolutionary distance in control conditions (Supplementary Figure 1B). We confirmed that *porB* and *rpmB* could diagnosis resistance phenotype after 10 minutes of ciprofloxacin exposure in a set of the isolates used in the prior study (Figure 4B,C). However, in evaluation of a broader diversity of the gonococcal population, the markers no longer showed diagnostic potential, displaying overlapping 2^−ΔΔCT^ values between resistant and susceptible isolates (Fig 4 B,C). These results suggest that the differential expression patterns of *porB* and *rpmB* are largely independent of resistance phenotype.

To our knowledge, our study is the first to comprehensively consider the genetic diversity of a species in development of RNA-based candidate diagnostic markers for resistance phenotype, and is also the first study to nominate RNA signatures diagnostic for azithromycin resistance in *N. gonorrhoeae*. Overall, our results emphasize the power and potential of these types of tests and also indicate the importance of selecting appropriate strains for the assay design panel in the research and development phase. We suggest that strain design panels must capture the genetic diversity of species for accurate evaluation of the factors underlying strain-to-strain gene expression heterogeneity, and control for the impact of phylogenetic structure on phenotypic diagnostics.

## METHODS

### Bacterial Strains and Culturing

*N. gonorrhoeae* isolates were acquired from the Center for Disease Control’s Gonococcal Isolate Surveillance Project (GISP). GISP isolates are collected from the first 25 cases of men with gonococcal urethritis per month at 25-30 sentinel sexually transmitted disease clinics across the United States (32). Isolates were cultured on GCB agar medium supplemented with 1% IsoVitaleX (Becton Dickinson Co., Franklin Lakes, N.J.). After inoculation, plates were incubated at 37°C in a 5% CO_2_ atmosphere incubator for 16-18 hours. All isolate stocks were stored at −80°C in trypticase soy broth containing 20% glycerol. Isolates for both azithromycin and ciprofloxacin test panels were selected to be representative of the known diversity within the gonococcal population. Antimicrobial susceptibility testing was conducted using E-test strips (40) or the agar dilution method (32), and MICs were recorded after 24 hours of growth. The European Committee on Antimicrobial Susceptibility Testing (EUCAST) defines azithromycin reduced susceptibility as MICs ≥ 1 μg/ml (41) while GISP uses a breakpoint of ≥ 2 μg/ml (32). Therefore, we conservatively define azithromycin reduced susceptibility in this study by the EUCAST standard of ≥ 1 μg/ml. For ciprofloxacin resistance we adhere to the GISP and CLSI breakpoint of ≥ 1 μg/ml (32, 42), which was also the breakpoint used by Khazaei et al. (3).

### Experimental Design

Cells harvested from overnight plates were suspended in GCP (7.5 g Protease peptone #3, 0.5 g soluble starch, 2 g dibasic K_2_HPO_4_, 0.5 g monobasic KH_2_PO_4_, 2.5 g NaCl, ddH_2_O to 500 ml; Becton Dickinson Co., Franklin Lakes, N.J.) supplemented with 1% IsoVitaleX and 0.042% sodium bicarbonate, and incubated at 37°C for 2 hours to mid-log phase. For azithromycin, samples were collected before exposure to azithromycin, 60 minutes post 0.125 μg/ml azithromycin exposure, and 180 minutes post 0.125 μg/ml azithromycin exposure (Figure 1). For ciprofloxacin we replicated the sampling scheme of Khazaei et al. (3), and sampled paired 0.5 μg/ml ciprofloxacin exposed and unexposed cultures 10 minutes post drug treatment (Supplementary Figure 3). RNA was isolated using the Direct-Zol kit with DNase I treatment (Zymo Research, Irvine, C.A.).

### RNA Preparation and Sequencing

RNA libraries were prepared at the Broad Institute at the Microbial ‘Omics Core using a modified version of the RNAtag-seq protocol (43). 500 ng of total RNA was fragmented, depleted of genomic DNA, dephosphorylated, and ligated to DNA adapters carrying 5’-AN_8_-3’ barcodes of known sequence with a 5’ phosphate and a 3’ blocking group. Barcoded RNAs were pooled and depleted of rRNA using the RiboZero rRNA depletion kit (Epicentre, Madision, W.I.). Pools of barcoded RNAs were converted to Illumina cDNA libraries in 2 main steps: (i) reverse transcription of the RNA using a primer designed to the constant region of the barcoded adaptor with addition of an adapter to the 3’ end of the cDNA by template switching using SMARTScribe (Clontech, Mountain View, C.A.) as described (44); (ii) PCR amplification using primers whose 5’ ends target the constant regions of the 3’ or 5’ adaptors and whose 3’ ends contain the full Illumina P5 or P7 sequences. cDNA libraries were sequenced on the Illumina Nextseq 500 platform to generate 50-bp paired end reads.

### RNA-seq Analysis

Barcode sequences were removed, and reads were then aligned to the FA1090 reference genome (AE004969.1) using BWA v.0.7.8 (45), mapping to the sense strand of the coding domain sequences (CDSs, n=1894). Differential expression analysis was conducted with DESeq2 v.1.10.1, which employs an empirical Bayes method to estimate gene-specific biological variation (46). A multi-factor analysis was used to account for the effects of resistance phenotype (*P*), condition (*C*), or the interaction between both phenotype and condition on transcript expression (*E*), using the model:

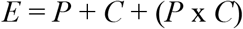

Per-gene dispersions were estimated using the Cox–Reid profile-adjusted method and a negative binomial generalized log-linear model was fit to identify significantly differentially expressed transcripts. Subsequently, gene lists were corrected for multiple testing using a false discovery rate (FDR) (cut-off of ≤0.01). We then compared significant model factors to define expression profiles for each transcript.

Genomic read libraries were obtained from the NCBI’s Short Read Archive (project PRJEB2090) and the European Nucleotide Archive (projects PRJEB2999 and PRJEB7904). Pairwise SNP distances were generated by mapping read libraries to the FA1090 (AE004969.1) reference genome using BWA v.0.7.8, and calling variants with Pilon v1.16 (47) using mapping qualities > 40 and coverage depths > 5x set as minimum thresholds. SNP distance between each isolate pair was then calculated excluding gaps using custom scripts. Normalized transcript read counts were generated for coding domain regions annotated in the FA1090 reference for each isolate (see above), and the absolute log-fold change was calculated by comparing transcriptional counts per CDS between each isolate pair at baseline (no drug exposure). CDS regions with zero counts in any isolate within each pair were excluded.

### RT-qPCR Validation

SuperScript IV reverse transcriptase (Thermo Fisher Scientific, Waltham, M.A.) was used to convert isolated RNA to cDNA using random hexamer primers. Annealing of primers to the template RNA was conducted in 13 μl reactions (3.8 μM primers, 1.3 mM dNTP mix, and 500 ng RNA) by heating the RNA-primer mix to 65 °C for 5 minutes and then incubating on ice for 1 minute. For reverse transcription, 4 μl of 5x SSIV Buffer, 1 μl of 100mM DTT, 1 μl of RNaseOUT, and 1 μl of SuperScript IV reverse transcriptase were added to the reaction mix and incubated at 23°C for 10 minutes, followed by 55°C for 10 minutes, and 80°C for 10 minutes.

Primers for azithromycin susceptibility diagnostic markers were designed to amplify ~100-bp fragments of both control and target genes (Supplementary Table 1). The control gene NGO1935 was empirically defined from our RNA-seq results, and was required to have no significant change in expression across treatments for all isolates tested. Primers for ciprofloxacin diagnostic markers were obtained from Khazaei et al. (3) (Supplementary Table 1). Real-time PCR was conducted with the Applied Biosystems PowerUp SYBR Green Master Mix on the Applied Biosystems ViiA 7 Real-Time PCR system (Thermo Fisher Scientific, Waltham, M.A.), using the following cycling conditions: 10 minutes at 95 °C, followed by 40 cycles of 95 °C for 15 seconds and 60-64 °C for 1 minute (see Supplementary Table 1 for primer-specific details). The 6.25 μl PCR reaction mix consisted of 1 μl of 1:100 diluted cDNA, 0.5x SYBR Green master mix, and 0.5 μM of the forward and reverse primer pair. Reactions were performed in 384-well plates with 3-4 biological replicates per isolate tested per antibiotic exposure condition. CT values were calculated using the ViiA 7 Software, and fold change was calculated using the 2^−ΔΔCT^ method (30, 31).

## ACKNOWLEDGEMENTS

We thank Dr. Jonathan Livny for his technical assistance with transcriptome library preparation, and Dr. Samantha Palace for her aid in RT-qPCR assay design.

## DATA ARCHIVING

Transcriptome libraries are archived in the GenBank SRA (Bioproject ID: PRJNA518111). RT-qPCR data are available in Supplementary Table 2.

## SUPPLEMENTARY FIGURES

**Supplementary Figure 1.**
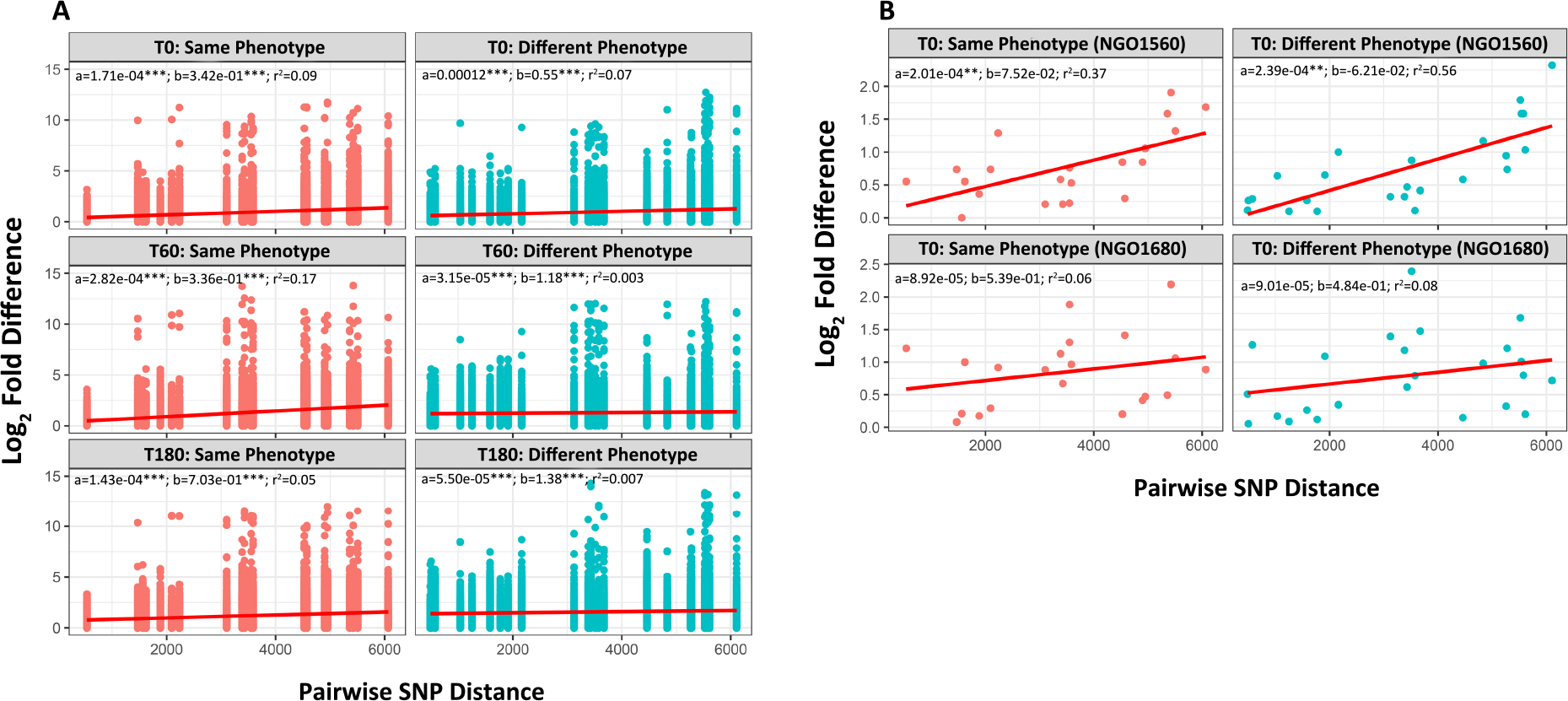
Transcriptome-wide differentially transcriptional regulation as a function of genetic distance in ten phylogenetically diverse isolates. **(A)** For all ten isolates for which we collected RNA-seq libraries, the number of pairwise SNP differences were compared to the per-transcript pairwise differences in gene expression at baseline (T0, no drug), after 60 minutes of drug exposure (T60), and after 180 minutes of drug exposure (T180). Regression lines were fit to either paired isolate comparisons that belonged to the same (resistant *vs.* resistant or susceptible *vs.* susceptible) or different (resistant *vs.* susceptible) phenotypic classes; and in all cases we observed positive and significant slope values. (B) For the candidate ciprofloxacin diagnostic *porB* (NGO1560) and *rpmB* (NGO1680) (32), there were generally stronger correlations and positive relationships.

**Supplementary Figure 2.**
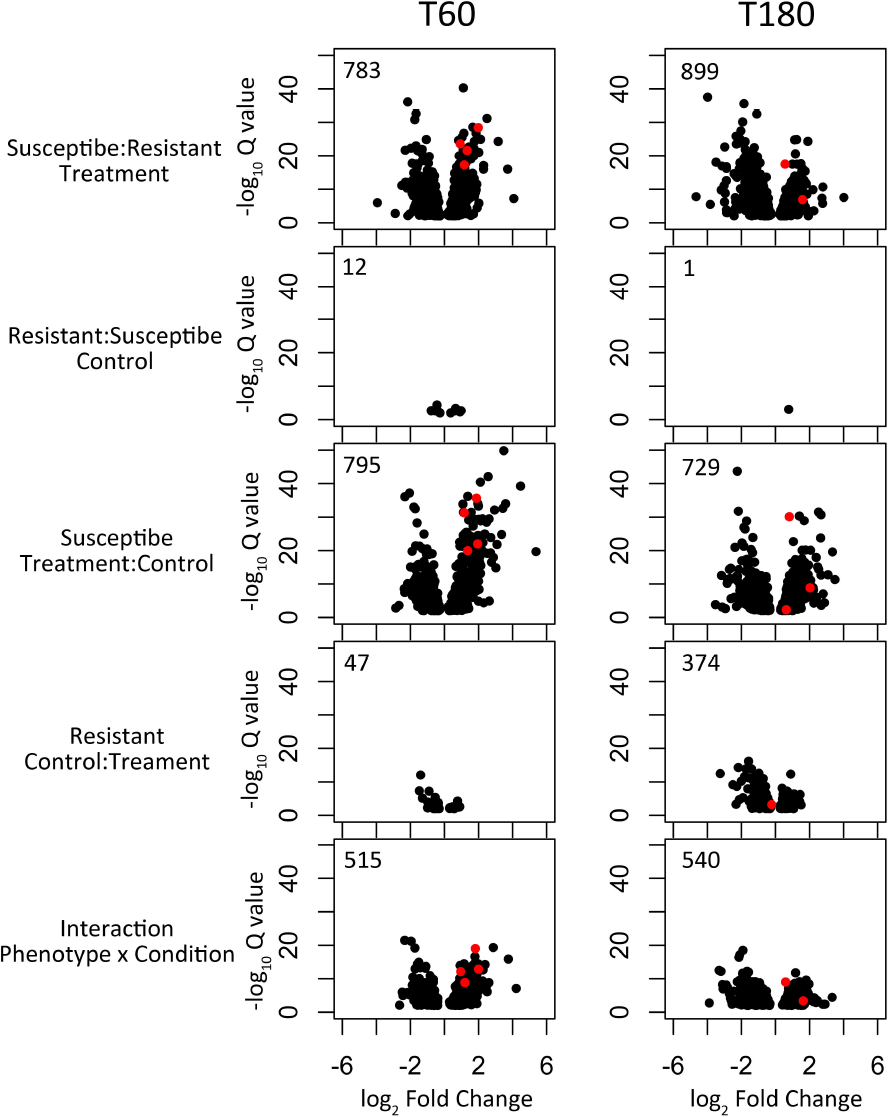
Volcano plots displaying the number of significantly differentially expressed transcripts (FDR cutoff ≤ 0.01) for each contrast of interest between the control to 60-minute and 180-minute azithromycin exposure conditions (black circles). The majority of transcripts were significantly differential expressed between resistant and susceptible isolates in the treatment conditions, and between the control and treatment conditions in susceptible isolates. Many transcripts also had a significant interaction term, suggesting that they had a different response trajectory between resistant and susceptible isolates across conditions. There was a high level of transcriptional response to azithromycin exposure in both the 60-minute and 180-minute treatments. Thus, diagnostic markers were selected from the earlier 60-minute exposure. These markers included: NGO0191, NGO1079, NGO1577, and NGO1829 (red circles) which all had highly significant model terms indicative of differential expression between phenotypes in the treatment condition, expression change across condition in susceptible isolates, and an interaction term.

**Supplementary Figure 3.**
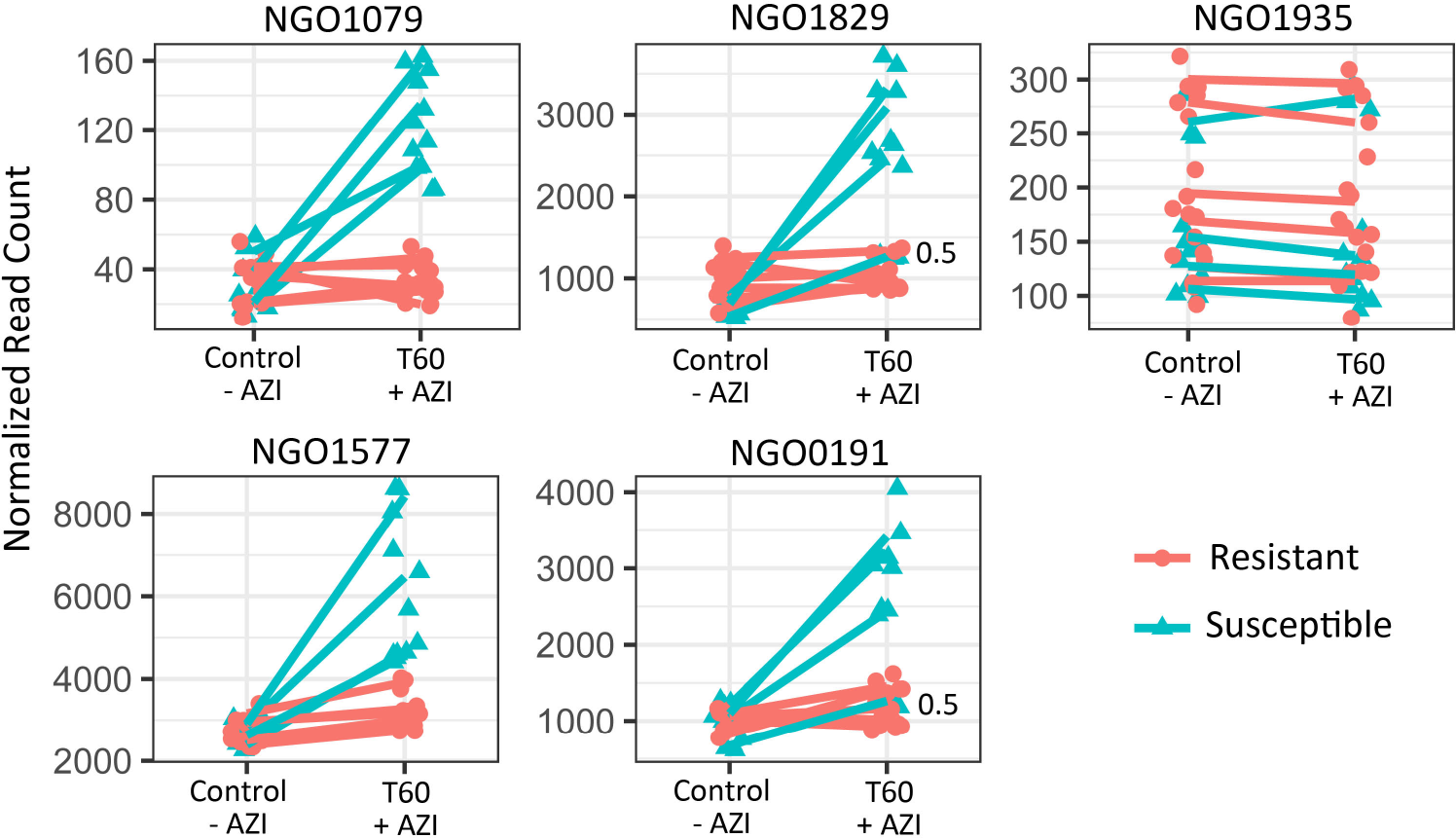
RNA-seq expression patterns for azithromycin diagnostic NGO0191, NGO1079, NGO1577, NGO1829 markers and the control NGO1935 marker. Read counts normalized by library size are plotted against the azithromycin exposure condition. The control condition was sampled prior to the addition of azithromycin to the culture media and the treatment condition was 60-minutes post exposure. Expression profiles across conditions are shown for the four susceptible isolates and six resistant isolates for which we collected RNA-seq data. All diagnostic markers had highly significant model terms indicative of differential expression between phenotypes in the treatment condition, expression change across condition in susceptible isolates, and an interaction term; though we did observe that isolates with intermediate MICS (i.e., 0.5 μg/ml) sometimes were more phenotypically similar to resistant or susceptible isolates dependent on the marker. The control NGO1935 gene was selected by the absence of significance in any of the tested glm contrasts.

**Supplementary Figure 4.**
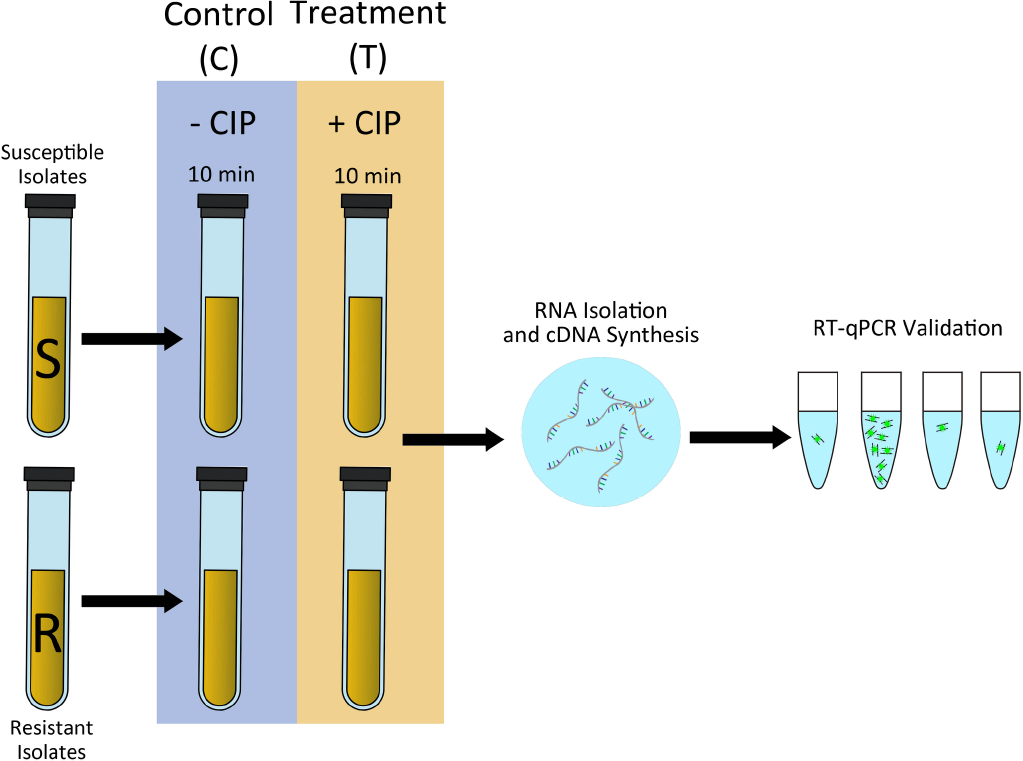
Workflow for validating ciprofloxacin resistance diagnostic markers. Susceptible (MIC < 1 μg/ml) and resistant (MIC ≥ 1 μg/ml) isolates were cultured in liquid GCP supplemented with 1% IsoVitaleX and 0.042% sodium bicarbonate. Paired cultures for each strain were either exposed to 0.5 μg/ml ciprofloxacin or unexposed, and then sampled after 10 minutes. RNA was isolated using the Direct-Zol kit and the SuperScript IV reverse transcriptase was used for first-strand cDNA synthesis. We then tested for differences in fold-change between condition across a panel of isolates using the nominated *porB* and *rpmB* (3) in reference to the control 16s rRNA gene.

**Supplementary Figure 5.**
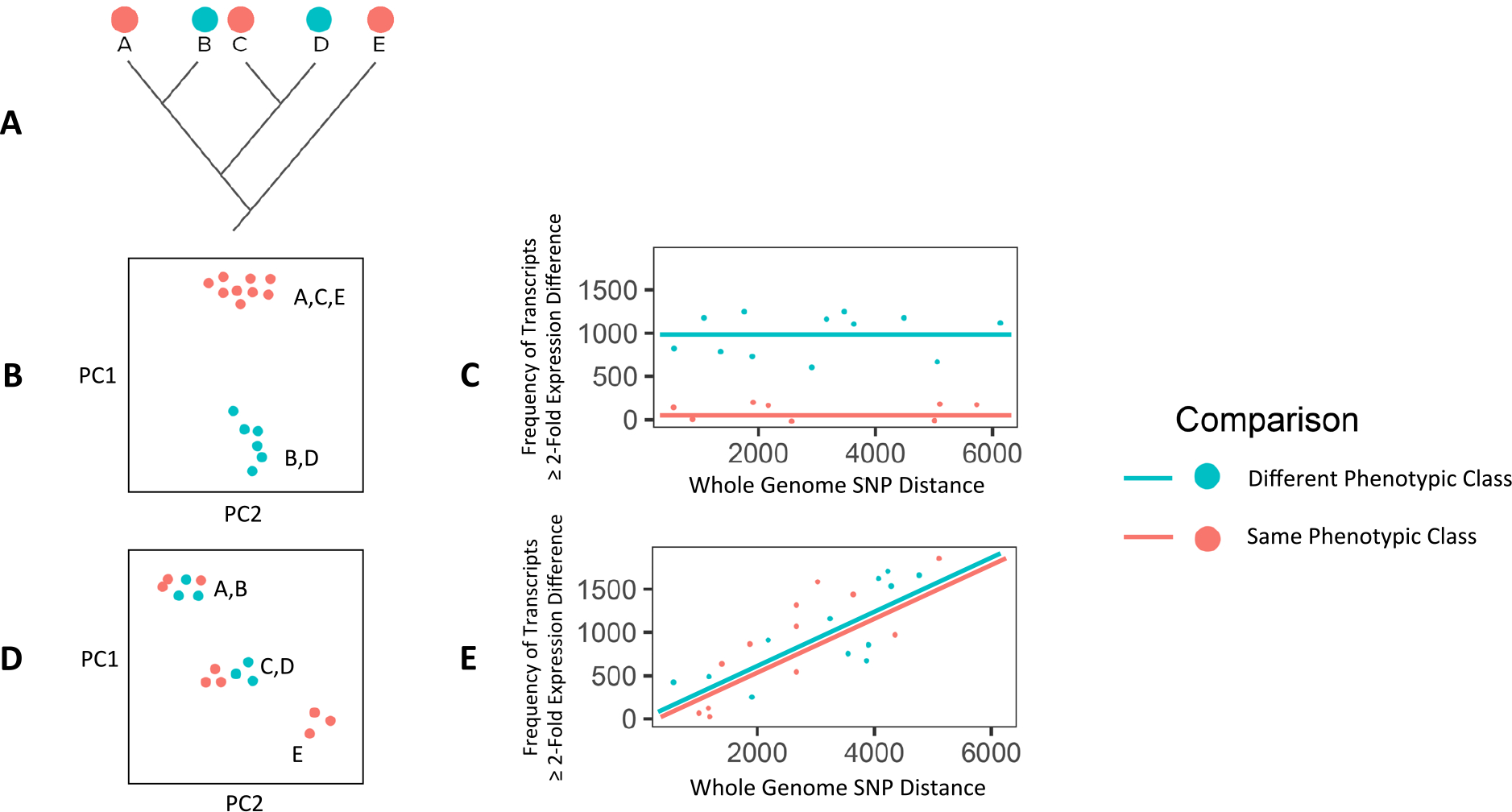
Contrasting the impacts of phenotypic class or genetic distance as the main dimension in driving transcriptional regulation. Here, we hypothesize that there are two major dimensions that may drive the majority of variance in transcriptional regulation between isolates: either phenotypic class (drug resistant *vs*. susceptible), or evolutionary time. (A) Given a phylogeny of a hypothetical population with branches indicating evolutionary distance and colors representing resistant (red) and susceptible isolates (blue): (B) if resistance phenotype is the major component driving transcriptional variation, isolates will transcriptionally cluster by phenotype as they are physiologically responding to the environment in the same way, and (C) regardless of the genetic distance of isolates, paired comparisons of isolates of the same phenotypes will show very little differences in transcriptional regulation, and paired comparisons of isolates of different phenotypes will display greater divergence in regulation of their transcriptomes. (D,E) If genetic distance is the major factor impacting transcriptional regulation, we expect that with increasing evolutionary time isolates will display increasing levels of differential transcriptome regulation.

